# Young Adults Recruit Similar Motor Modules Across Walking, Turning, and Chair Transfers

**DOI:** 10.1101/2021.05.04.442677

**Authors:** Hannah D. Carey, Daniel Liss, Jessica L. Allen

## Abstract

Moving about in the world during daily life requires executing and successfully shifting between a variety of functional tasks, such as arising from a chair or bed, walking, turning, and navigating stairs, etc. Moreover, moving about during daily life requires not only navigating between different functional tasks but also performing these tasks in the presence of mental distractions. However, little is known about underlying neuromuscular control for executing and shifting between these different tasks. In this study, we investigated muscle coordination across walking, turning, and chair transfers by applying motor module (aka muscle synergy) analysis to the Timed-Up-and-Go (TUG) test with and without a secondary cognitive dual task. We found that healthy young adults recruit a small set of common motor modules across the subtasks of the TUG test and that their composition is robust to cognitive distraction. Instead, cognitive distraction impacted motor module activation timings such that they became more consistent. This work is the first to demonstrate motor module generalization across multiple tasks that are both functionally different and crucial for healthy mobility. Overall, our results suggest that the central nervous system may draw from a “library” of modular control strategies to navigate the variety of movements and cognitive demands required of daily life.

**New & Noteworthy:** We demonstrated that healthy young adults recruit a small set of motor modules across subtasks of the Timed-Up-and-Go test (i.e., walking, turning, and chair transfers). Moreover, we showed that motor module composition, but not activation timing, is robust to cognitive distractions. These results support the hypothesis that healthy young adults recruit from a “library” of motor modules and modulate their activation timing to meet the different mechanical and cognitive demands required to navigate daily life.

## 1. Introduction

Moving about in the world during daily life requires executing and successfully shifting between a variety of functional tasks, such as arising from a chair or bed, walking, turning, and navigating stairs, etc. This multi-task nature of daily life is recognized clinically, with many clinical tests of mobility assessing multiple functional tasks (e.g. Timed-Up-and-Go test (1), Berg Balance test (2), Mini BESTest (3)). In contrast, the neuromuscular control underlying the execution of and coordination between different functional tasks is less understood. Although the neuromuscular control of different functional tasks have been studied in isolation (e.g., locomotion (4), standing (5), etc.), little is known about how neuromuscular control compares *across* different functional tasks. A better understanding neuromuscular control across different functional tasks will provide valuable insight into the strategies that enable us to successfully navigate the many tasks required for daily life.

Motor module analysis is commonly used to investigate neuromuscular control strategies during movement (e.g., (6–14) etc.). Motor modules, or muscle synergies, are groups of coactive muscles flexibly recruited over time to meet the biomechanical demands required of a functional task (15). To date, motor module analysis has primarily been used to investigate neuromuscular control within a single functional task. Such studies provide evidence that similar motor modules are recruited within the same functional task under different task demands, such as level versus inclined running (16), varied pedaling speeds (17), straight versus curved walking (18), and reactive balance during different stance positions (19). In each case, changing musculoskeletal configurations or mechanical demands were addressed with changes in temporal activation and/or incorporation of task-specific motor modules rather than a new set of modules for each condition. While this implies that the nervous system may rely on a common set of motor modules to accomplish a variety of conditions for a particular task, we do not know whether this motor module generalizability extends to a broader range of functionally different tasks.

Motor module generalization, or recruiting common motor modules across functionally different tasks, may enable the successful execution and switching between tasks. Initial evidence for motor module generalization comes from animal studies, where, for example, frogs were found to recruit common motor modules across walking, swimming, and jumping tasks (20). Although seemingly all locomotive tasks, the joint mechanics required to produce them are different in each task. More recently, evidence that such motor module generalization also occurs in humans has emerged. In particular, we recently found that young adults recruit common motor modules across standing reactive balance and unperturbed walking (21) and that reduced generalization across these two functionally different tasks was associated with impaired gait, balance, and mobility performance in both neurotypical and neurologically impaired populations (e.g. young adults (21), stroke (22), PD (23)). Such a relationship provides support for motor module generalization as a neuromuscular control strategy for successful mobility during daily life. However, the extent to which motor modules are generalized across the wider range of functional tasks encountered during daily life (e.g., walking, turning, chair transfers, etc.) remains unclear.

Moving about during daily life requires not only navigating between different functional tasks but also performing these tasks in the presence of mental distractions (i.e., cognitive-motor dual tasking). Putting on a jacket while carrying on a conversation or walking through store aisles while trying to remember the items on a grocery list are common examples of cognitive-motor dual tasking in everyday life. Because the biomechanical requirements of any functional task are the same with or without cognitive distraction, it is likely that the same motor modules are recruited in both scenarios. Instead, the mental distraction may pull away some of the cognitive resources normally used to plan and generate movement, muddying the typical command signals and leading to changes in temporal motor module activation. It is known that dual-task conditions result in increased variability in gait parameters (e.g. stride time (24), or swing time (25) in older adults) but the impacts on muscle activation or motor module recruitment is not well characterized. However, increased gait variability suggests that the motor module activations producing gait may also become more variable (e.g., from step to step during walking). Identifying the differences in motor module recruitment between distracted and undistracted tasks may provide valuable insight into neuromuscular control strategies for achieving common daily tasks.

In the present study, we analyzed electromyography (EMG) collected from the hip, knee, and ankle muscles from young adults while performing the Timed-Up-and-Go (TUG) test to investigate motor module generalization across different functional tasks. The TUG test is a commonly used clinical mobility test in which subjects stand up from a chair, walk 3 meters, turn around a cone, and walk back to the chair to sit down (1). We chose to examine muscle activity during the TUG test because it contains a variety of functional tasks that are important for daily life. In particular, the TUG test includes transitional subtasks like chair transfers and turns that are critical for independence but also a common source of falls (26–28). Our overall hypothesis is that healthy young adults recruit from a “library” of motor modules to meet the multi-task demands of daily life and that motor module composition is robust to cognitive distractions. Based on this hypothesis, we predicted that (1) young adults would recruit a small number of common motor modules across the subtasks of the TUG test (sit-to-stand, walking, turning, and stand-to-sit) and that when performing a secondary cognitive task (2) the number and composition of these motor modules would not change, (3) but their recruitment timing and level of activation would become more variable.

## 2. Methods

### 2.1 Participants

Thirteen healthy young adults (5 M, 21.4±1.6 yrs) participated in this study. Inclusion criteria was age between 18-35 years old. Exclusion criteria were any diagnosed neurological or psychological conditions, musculoskeletal conditions, sensory deficits, stroke, traumatic brain injury, or a concussion or other injury within a year of participation. All participants provided written informed consent before participating according to an experimental protocol approved by the institutional review board of West Virginia University.

### 2.2 Data collection and processing

Each subject performed the TUG test first while walking normally (TUG) and then while counting backwards by three’s (TUGC). For TUGC, subjects were instructed to pay equal attention to both the counting and walking tasks. Subjects self-selected which direction they turned around the cone until 10 trials of one turn direction were completed. Then we instructed them to turn the opposite direction for the remaining trials. Turning direction when sitting back down in the chair was not enforced. Some trials were removed before analyzing due to experimental or equipment error (n=25, 5% of total trials) or subject error (e.g., kicking the cone, n=21, 4% of total trials). In both conditions, each subject completed the TUG test with at least 6 good trials for each turn direction around the cone (avg: TUG 9.46±1.42, TUGC 10.12±1.30).

Three-dimensional marker position was collected at 100 Hz with a 10 camera Vicon motion capture system and a modified plug-in gait marker set with 31 markers placed on the head, trunk, pelvis, thigh, shank, and foot segments. Marker data from the heels, toes, and clavicle were used to segment the TUG test into 4 subtasks: Sit-to-Stand, Walk, Turn, and Stand-to-Sit. The two walking portions were combined into one subtask and turn directions for both the Turning and Stand-to-Sit subtasks were considered separately (e.g., right turn vs. left turn) for a maximum total of 6 subtasks. Turning direction during Stand-to-Sit was not enforced; some subjects consistently chose one direction for every trial and therefore only had 5 different subtasks. Details of TUG segmentation are listed in Table 1 and an example can be found in the supplementary material (Figure S1).

Surface EMG data were collected at 1000 Hz from 12 muscles per leg: gluteus maximus (GMAX), gluteus medius (GMED), tensor fasciae latae (TFL), adductor magnus (ADD), biceps femoris long head (BFLH), rectus femoris (RFEM), vastus lateralis (VLAT), medial and lateral gastrocnemius (MGAS and LGAS), soleus (SOL), peroneus (PERO), and tibialis anterior (TA). EMG data were high-pass filtered at 35 Hz, demeaned, rectified, and low-pass filtered at 10 Hz using custom MATLAB scripts (example EMG in TUG, Fig. 1B). For each subject, leg, and condition, separate EMG matrices were generated by concatenating data from all trials for that condition. Or in other words, for each condition (TUG and TUGC), there were 6 or 7 different EMG matrices per subject and leg (each subtask plus the full TUG test). Those subjects who consistently turned in the same direction when sitting back down had 6 matrices, whereas those who mixed their turning direction when sitting down had 7. Each EMG matrix was then normalized to the maximum observed value for each muscle in the EMG matrix for the full TUG test.

**Figure 1:**
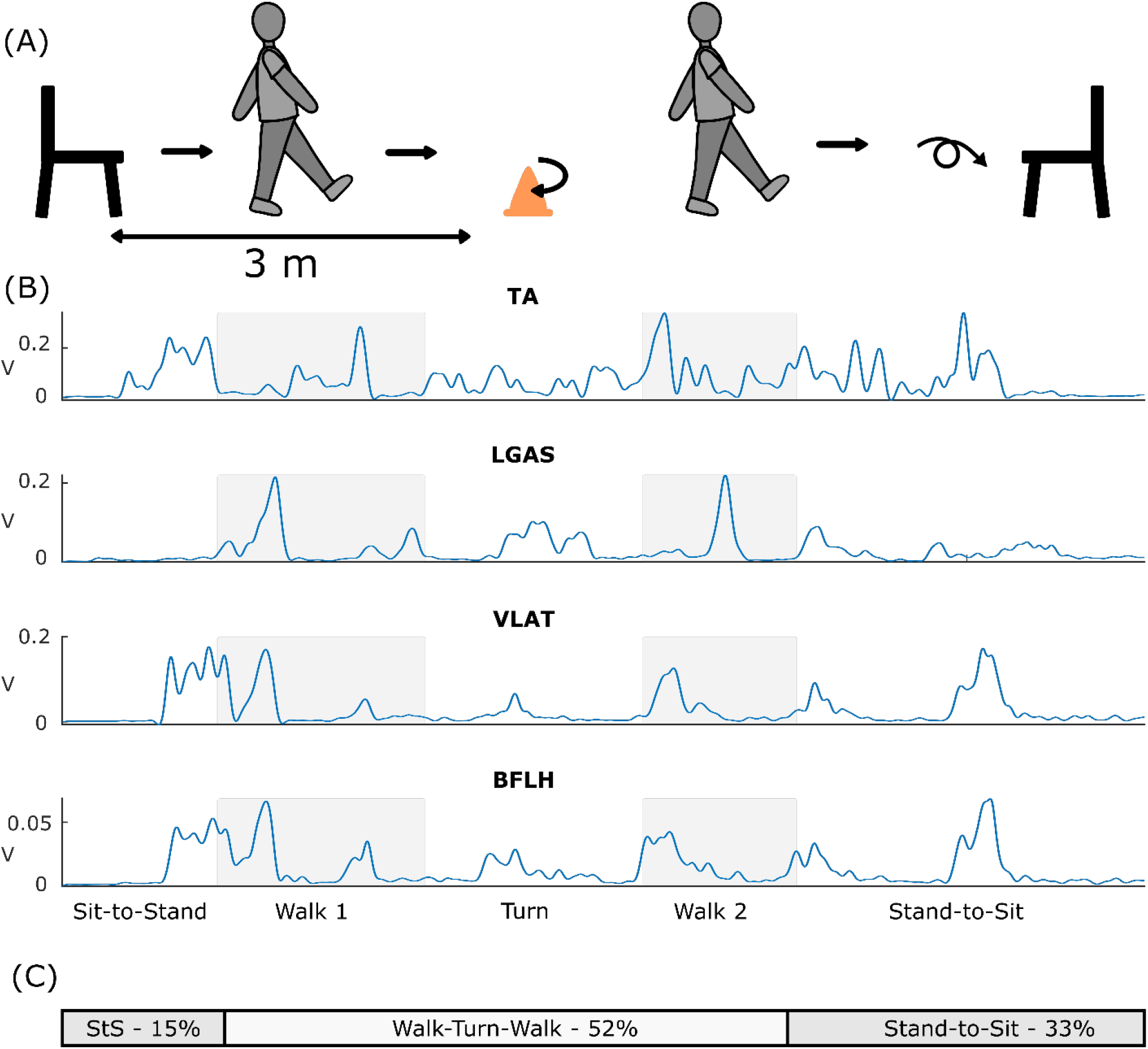
The Timed-Up-and-Go (TUG) test. (A) In the TUG test, subjects get up from a chair, walk around a cone 3 m away, walk back to the chair, and sit back down. (B) Example muscle activity from selected muscles (tibialis anterior (TA), lateral gastrocnemius (LGAS), vastus lateralis (VLAT), and biceps femoris long head (BFLH)) during the TUG test with labeled subtasks. Gray boxes indicate the walking portions of TUG, while white sections indicate Sit-to-Stand, Turn, and Stand-to-Sit. (C) The subtask proportions used during activation analyses (see section 2.3.2).

### 2.3 Motor modules extraction and analysis

Motor modules were separately extracted for each subject, leg (left vs. right), and condition (TUG vs. TUGC) using non-negative matrix factorization from each EMG data matrix. (i.e., the full TUG test and each TUG subtask). Motor modules were extracted such that EMG=W x C + ∊, where W is an *m* x *n* matrix of the *n* motor module weights for *m* muscles, C is an *n* x time matrix containing the activation coefficients for each module, and ε is the EMG reconstruction error. To ensure equal weight of each muscle during the extraction process, the data for each muscle were scaled to unit variance before motor module extraction and then rescaled to original units afterwards.

We extracted 1-12 motor modules from each EMG matrix and selected the minimum number needed to sufficiently reconstruct the original data. Module numbers were chosen such that the 95% confidence interval of the overall variance accounted for (VAF) was greater than 90%, where VAF is the squared uncentered Pearson’s correlation coefficient between the reconstructed EMG (W x C) and the original EMG (21). 95% confidence intervals on the VAF were generated by resampling the original EMG 250 times with replacement and calculating the VAF for each sample. We then examined motor module generalization and the impact of the cognitive task as follows:

#### 2.3.1 Generalization of motor modules across tasks

To investigate motor module generalization during the TUG test, we used a clustering analysis to group similar modules recruited during the TUG subtasks. For each subject we determined (1) the level of motor module generalization across TUG subtasks, (2) the level of similarity between clustered motor modules, and (3) the level of similarity between modules recruited during TUG subtasks to those recruited during the full TUG test. Examples of these metrics are shown in Figure 2 and their calculations are described below.

**Figure 2:**
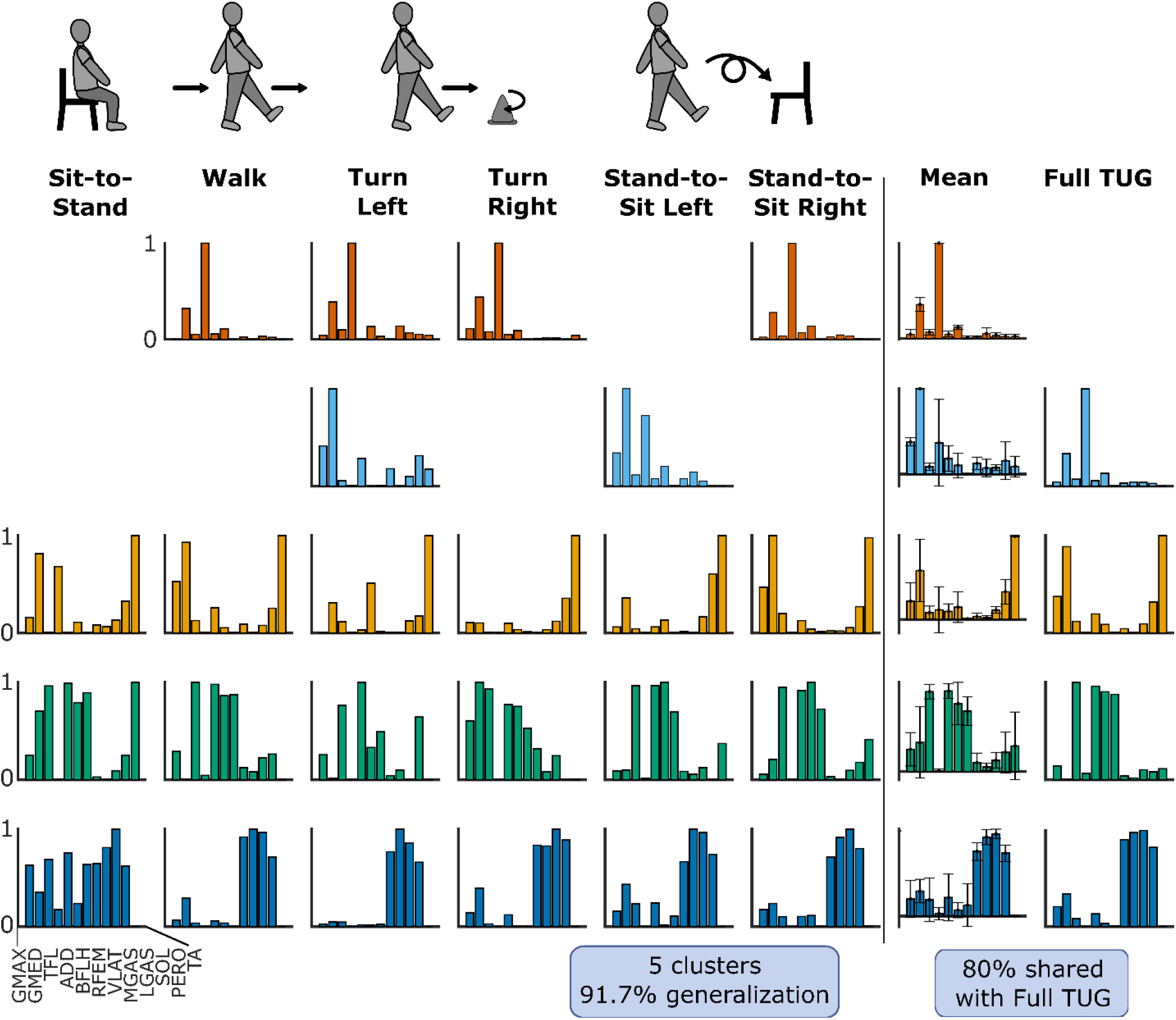
Example motor module clusters. Example of clustered motor modules for a representative subject’s left leg. The first 6 columns contain the motor modules recruited during each TUG subtask. Modules in the same row were clustered together. The second column from the right shows the average modules for each cluster and the last column contains the motor modules from the full TUG test. In this example, the subject had 5 clusters and 91.6% generalization. There are 4 common motor modules between the full TUG test and the cluster averages, giving 80% in common.

1. <u>Motor module generalization.</u> Motor modules recruited during TUG subtasks in the normal condition were separately pooled for each subject and leg and then sorted with a hierarchical clustering algorithm (7). The ‘cluster’ function from the MATLAB Statistics and Machine Learning Toolbox was used to cluster the modules, with the distance metric Minkowski order p=3 and Ward’s linkage option. The number of clusters within each group was determined as the minimum number such that each cluster contained no more than one motor module from each subtask (7,8,22). Generalization of motor modules across subtasks was calculated as a percentage and defined as,

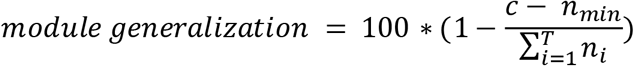

where *c* is the number of clusters, *n_i_* is the number of modules recruited during the *ith* subtask, T is the total number of subtasks (5 or 6 per subject, depending on whether a subject turn in both directions in the Stand-to-Sit turns or not), and *n_min_* is the smallest number of modules recruited in that subject and leg during any subtask.
2. <u>Within-cluster motor module similarity.</u> To assess module similarity within each cluster, we calculated the cluster consistency as the pairwise linear correlation coefficient between all modules in each cluster and averaged for each subject and leg. Module pairs with r≥0.7079, the critical r^2^ value for 12 muscles at α=0.01, were considered similar.
3. <u>Similarity between sub-task and full TUG motor modules.</u> Finally, to determine the similarity of modules identified during the TUG subtasks to modules from the full TUG test, motor modules from the full TUG test were compared to averaged modules from each cluster using Pearson’s correlation coefficients, again with a similarity threshold of r≥0.7079.

#### 2.3.2 Effects of a cognitive task on motor module recruitment

To characterize the effects of a secondary cognitive task on motor module recruitment, we compared both the spatial and temporal aspects of motor modules recruited during TUG versus TUGC.

We analyzed spatial effects by comparing (1) motor module number and (2) motor module composition between TUG and TUGC. The number of motor modules recruited during TUG and TUGC were compared using paired t-tests for the full TUG test and each of its subtasks (7 total). Motor module composition (W’s) from TUG and TUGC for the full TUG and each subtask were compared using Pearson’s correlation coefficients, where module pairs with correlation coefficients r≥0.7079 were considered the same. We also identified how many modules were common between TUG and TUGC by calculating the percentage of common modules, defined as,

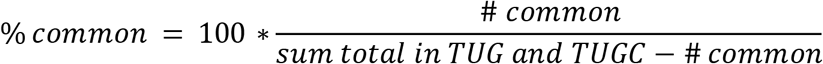

for each subject, leg, and subtask.

We analyzed temporal effects by comparing motor module recruitment variability between TUG and TUGC. Motor module activation coefficients (C’s) for each synergy were first separated by trial. Each trial was then time-normalized to be the same number of data points and such that the lengths of the chair transfers and walking-turning portions were consistent. Specifically, for each trial we calculated the proportion of each segment as *subtask time / TUG time*. We then averaged these values across all trials and subjects and rounded to the nearest whole number for each TUG segment (Fig.1C.). Each trial was then normalized to be 1024 points long, with 154 data points in sit-to-stand, 532 points in walking-turning, and 338 points in stand-to-sit. See Fig S2 and “Normalization of Motor Module Activations” in the Supplementary Material for an example and further details. We then separated the trials based on “kinematic strategy”, defined as the sequence of first step leg, turn direction, and Stand-to-Sit turn direction. We separated trials in this way because the shapes of motor module activation curves vary based on the TUG kinematic strategy used (e.g., which leg was used to take the first step) without representing true changes in motor module recruitment. To account for this, we only compared the time-normalized module activations from sequences that a subject used in both TUG and TUGC. Specifically, the average root-mean-square error (RMSE) of module activations from common motor modules across all subjects, legs, and tasks were compared using a paired t-test. See Fig S3, Table S2, and “Kinematic Strategy Separation” in the Supplementary Material for an example and further details.

#### 2.3.3 Effects of dual task on TUG and counting performance

Finally, to investigate dual-task effects on cognitive performance (i.e., counting backwards by threes from a random number), we compared the counting score and counting rate during TUGC to baseline counting performance. Baseline counting performance was collected while subjects were seated in the chair for 15 seconds (minimum 2 baseline trials). Subjects were instructed to repeat the given number and then for each TUGC trial, the <u>counting score</u> was calculated as *# correct / total # of counts* and the <u>counting rate</u> as *total # of counts / time*. Counting scores during both TUGC and the baseline were highly skewed towards 1 (Shapiro-Wilk (sw) test statistics: baseline sw=0.50, p<0.001, TUGC sw=0.82, p=0.01), so they were compared using a Wilcoxon signed rank test (α=0.025). Counting rates during TUGC and the baseline fit within a normal distribution and were compared using a paired t-test (baseline sw=0.97, p=0.88, TUG sw=0.92, p=0.22). TUG performance times with and without the cognitive task were compared using a paired t-test.

## 3. Results

Subjects recruited a small number of unique modules that were similar across TUG subtasks. Motor modules from TUG subtasks were grouped into a small number of clusters (avg 5.6±0.99, Fig 3A), leading to a high percentage generalization (avg 89.23±3.41%, Fig 3B). Most clusters were consistent across subtasks (avg 0.80±0.06, Fig 3D), with only two of the 11 subjects having an average cluster consistency below the 0.7079 similarity threshold in one of their legs (avgs for each subject: 0.60,0.70). The averaged motor modules across all subtasks within each cluster were very similar to modules recruited during the full TUG test (avg r=0.789±0.115, Fig 3C). Motor module composition was unchanged when performing the TUG test with the secondary cognitive task of counting backwards by threes. Subjects recruited an average of 4.5 synergies during TUG (Fig 4), which was not significantly different during TUGC (p=0.75, Fig 5A and Supplementary Data Table 3). Similarly, there was no significant difference in the number of motor modules recruited during TUG and TUGC for any TUG subtask (see Supplementary Data Table 3 for all t-test results). Subjects recruited motor modules with similar compositions during TUG vs TUGC. Motor modules were highly similar during full TUG (93.7±0.1%, Fig 5A). Modules were also similar in each subtask (avg across all subtasks: 78.7±0.2), though there was more inter-subject variability (range = 17-100%, Fig 5B). Further, most module pairs were more strongly correlated than the similarity threshold, illustrated in a histogram of pooled correlation coefficients (Fig 5C).

**Figure 3:**
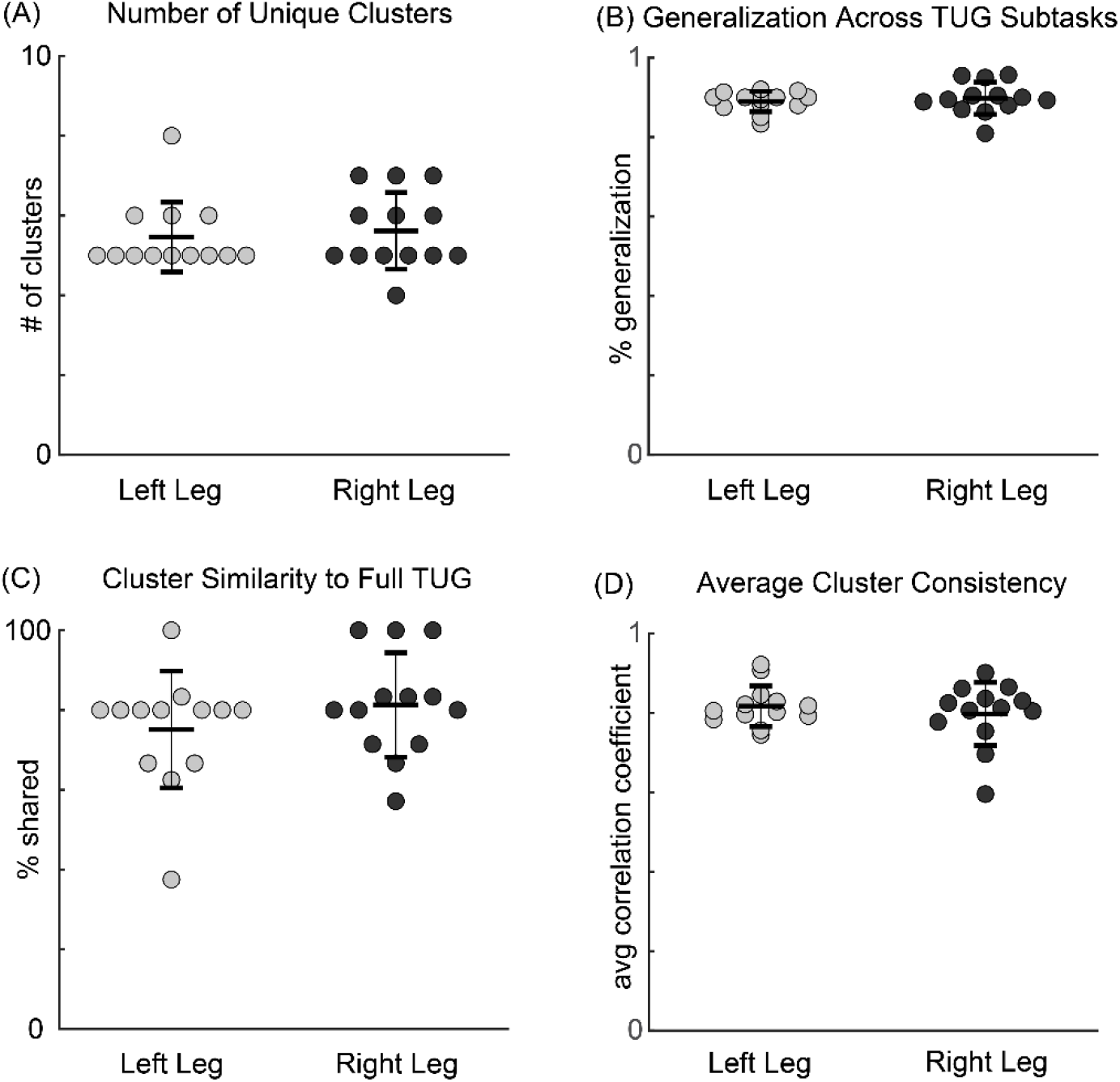
Motor module clustering results. For all panels, each dot represents one subject and leg (n=13). (A) Motor modules were grouped into a small number of clusters across all subjects, (B) leading to a high percentage generalization. (C) Motor modules recruited during the full TUG test were well matched with the cluster averages and (D) Motor modules within each cluster were similar to each other, producing a high cluster consistency.

**Figure 4:**
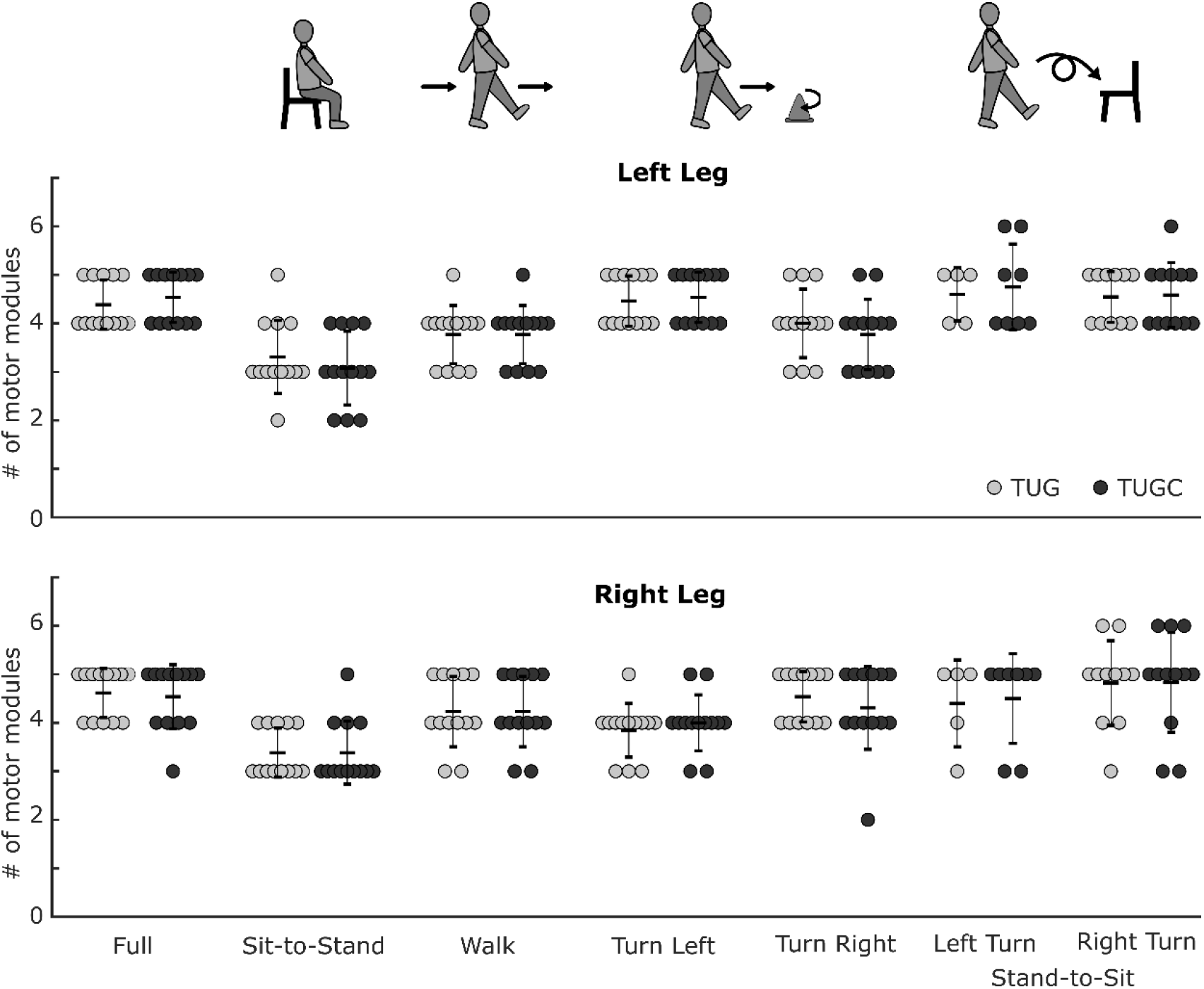
Number of motor modules recruited during the TUG test and its subtasks. The number of modules didn’t change between TUG and TUGC for the full TUG test, or any of the subtasks. (n=13, paired t-test p=0.75)

**Figure 5:**
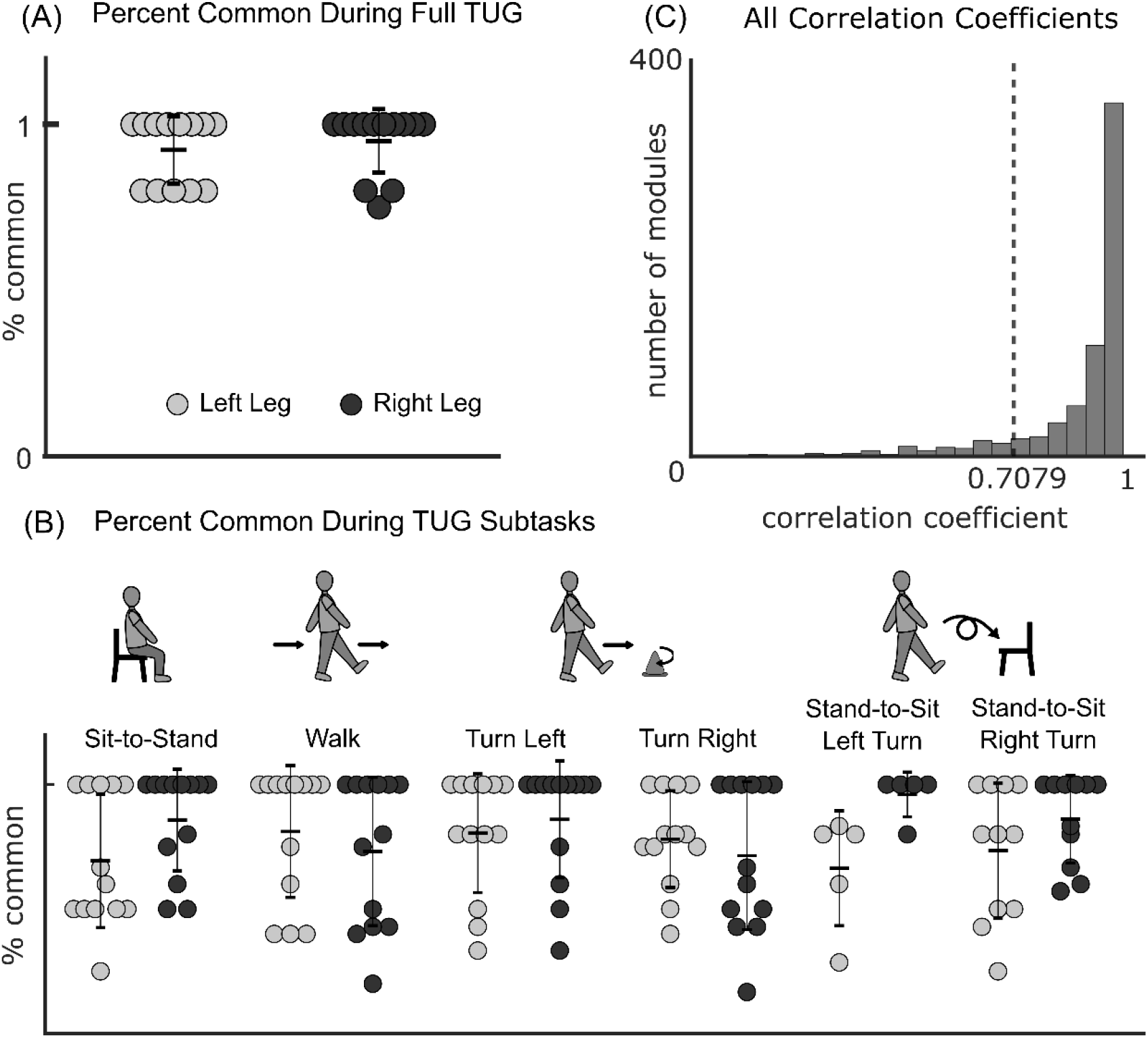
Motor module similarity during TUG and TUGC. (n=13) Motor module composition was very similar during TUG and TUGC, leading to a high percentage common during (A) the full TUG test and (B) each of its subtasks and (C) very high correlation coefficients between all pairs of modules (pooled across all subjects and subtasks, the gray line represents the cutoff for significant similarity, r≥0.7079).

In contrast, motor module activation became more consistent across repetitions of the TUG test when counting backwards by threes. Motor module activation variability was significantly lower in TUGC than in TUG (avg rmse for TUG: 0.066±0.010, TUGC: 0.061±0.011, p=0.008, Fig 6B).

**Figure 6:**
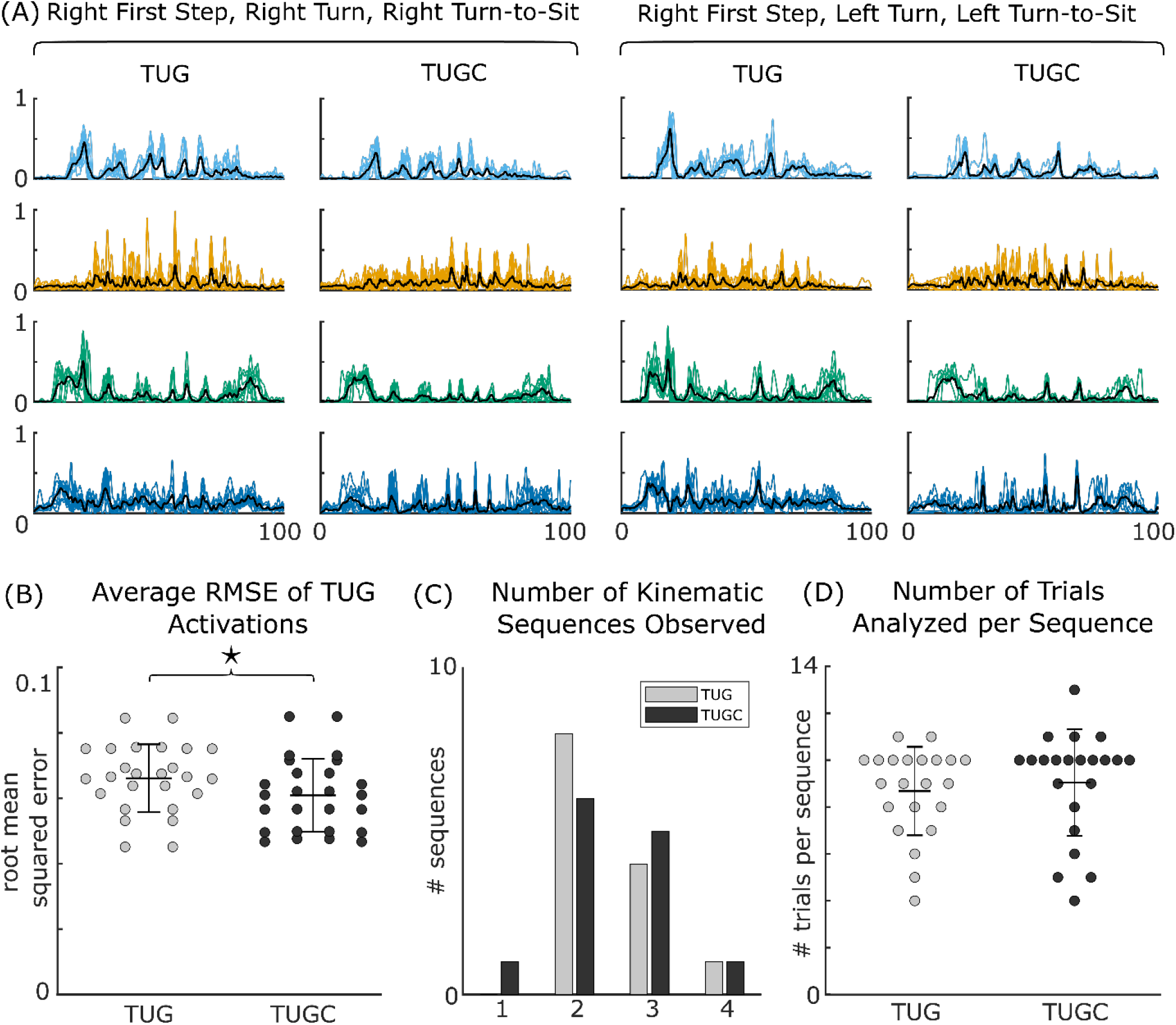
Temporal dual task effects. (A) Example module activations from the left leg of one subject in two kinematic strategies. (B) Average root mean squared error of motor module activations during TUG and TUGC (n=26 legs, paired t-test p=0.008). Module variability was significantly lower in TUGC than normal TUG. (C) Number of kinematic strategies (sequences) used by each subject. Across all trials, most subjects used 2-3 different kinematic strategies, but only had 1-2 strategies used in both TUG and TUGC. (D) Number of trials used in RMSE analysis, ranged from 4-13 trials per kinematic sequence

Importantly, the shape of the motor module activation curves varied depending on which leg took the first step, the turn direction, around the cone, and the turn direction when sitting back down (e.g., Fig 6A). Although most subjects used only two sequences (one for each turn direction around the cone), a smaller subset used 3-4 (Fig 6C) because they switched their turn direction when sitting down or varied the first step leg. Only the module activations from trials with similar sequences were compared between TUG and TUGC (avg 8.9±2.1 trials per sequence; Fig 6D).

Dual task affected TUG time but not counting performance. The addition of a secondary cognitive task led to a significant but small difference in TUG performance time (TUG: 6.76±0.93 s, TUGC: 7.11±1.10 s, p=0.02, Fig 7A). Counting score (base: 0.93±0.13, TUGC: 0.93±0.20, p=0.23, Fig 7B) and counting rate (base: 0.6590±0.24, TUGC: 0.0.63±0.16 counts/s, p=0.31, Fig 7C) was not different between the baseline trial and TUGC.

**Figure 7:**
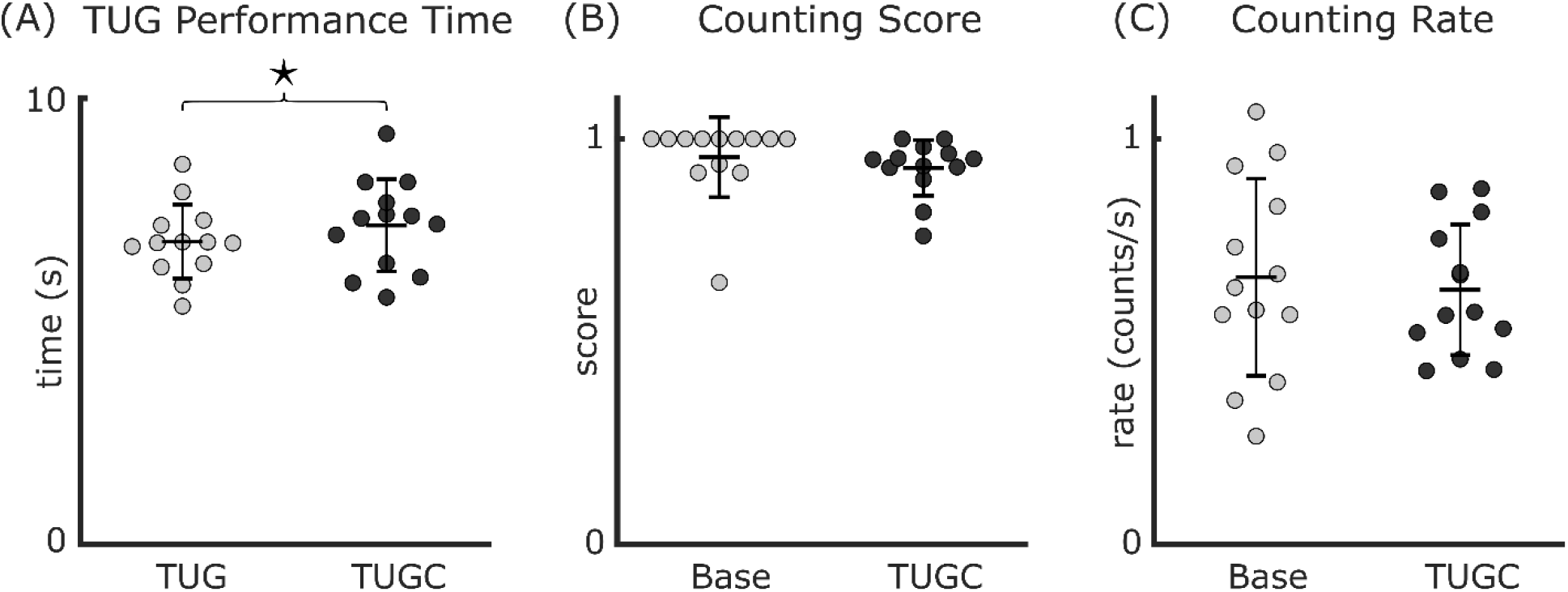
Dual Task Costs of the TUG test. (A) There was a small but significant increase in TUG performance time with the added counting task (n=13, paired t-test p=0.02). There was no change in either (B) counting accuracy (n=13, Wilcoxon signed rank test p=0.23) or (C) the counting speed (n=13, paired t-test p=0.22) from baseline to TUGC.

## 4. Discussion

The central nervous system may rely on generalizable control strategies to meet the multi-task demands of daily life. In support of this hypothesis, we show here that healthy young adults recruit a small set of generalizable motor modules across the subtasks of the TUG test and that the composition, but not the activation, of those motor modules is robust to cognitive distraction. This work is the first to demonstrate motor module generalization across multiple tasks that are both functionally different and crucial for healthy mobility.

### 4.1 Generalization of motor modules across tasks

Consistent with our hypothesis, our data suggest that young adults recruit a small set of generalizable motor modules across several functional tasks important for moving about in the world during daily life - walking, turning, and getting in and out of a chair. Prior work has demonstrated that the same motor modules are recruited to perform a single task under varying demands (e.g., pedaling at different speeds (17) or maintaining balance under different postural configurations (19)). Here, we expand upon this prior work to demonstrate that many of the same motor modules are recruited to perform *different* tasks.

Generalizing the recruitment of motor modules may enable the successful execution of similar basic mechanical demands required of different tasks. Except for turning, the tasks we examined are dominated by sagittal plane motion that likely require the achievement of similar basic mechanical demands such as plantarflexion, leg support, and center-of-mass stabilization. Even though our 180° turning task includes substantial non-sagittal plane motion, its successful performance also requires the achievement of many of these same demands. However, how these demands must be met and coordinated together to achieve successful task performance varies between tasks. For example, walking and sit-to-stand both involve propelling the center of mass forward and extending the limbs while keeping the foot fixed; however, sit-to-stand uses symmetric movements and includes a larger vertical COM movement, while walking alternates leg movements and requires stability during single leg stance (29,30). To meet these varying coordination requirements, we found that young adults modulated the recruitment (i.e., activation timing) and not the structure of the motor modules. We also found that most subjects recruited a plantarflexor module, knee extensor module, and a dorsiflexor module across all tasks. These motor modules are similar to those previously identified as important for meeting the mechanical demands of walking (31–33). Follow-up studies are needed to determine whether these generalized motor modules are indeed recruited to produce similar basic mechanical demands across different tasks.

Although many motor modules were generalized across all tasks, task-specific modules did emerge during turning. The emergence of task-specific modules is consistent with prior work. For example, Ivanenko and colleagues observed the emergence of task-specific modules when walking while performing an additional task (e.g., picking up an object or stepping over an obstacle) (34). However, the emergence of *turning*-specific motor modules differs from a study by Chia Bejarano and colleagues in which similar motor modules were recruiting during walking and turning (18). The contrasting results likely stem from differences in the differing radii of the turns and the mechanical demands they require. In (18), subjects walked around a circle with a 1.2 m radius, whereas in the current study subjects turn tightly around a cone or pivot on one leg to change direction 180° (see the left turns in Fig. 2). Such a tight turn may involve much more weight shifting and stepping changes than walking around a wider curve, and therefore are more likely to require additional motor module recruitment. For example, the inside turn leg would have increased demand for both stability and directing the turn. In our study, the turning-specific modules were often composed primarily of hip muscles (GMAX, GMED, ADD); GMED specifically is known to be important for pelvic stability during single leg stance (35,36)and contributes to mediolateral control of the center-of-mass (37). The recruitment of such a module is consistent with increased demand for stability and frontal plane movements during this turn that may not be achievable using the generalized modules on their own. As turns are a common source of falls for people with mobility impairments (e.g., (26,27,38)); some of this difficulty could stem from an inability to appropriately recruit turning-specific motor modules. Overall, our results suggest that the nervous system reuses and modifies the same control strategies to execute and shift between similar tasks. When the mechanical demands for a task cannot be met by that module set, additional modules must be recruited.

### 4.2 Dual Task Effects

Consistent with our hypothesis, we found that motor module number and composition are robust to cognitive distraction. Moreover, we found that both TUG and counting performance were not affected by the cognitive-motor dual task condition. Though we identified a statistically significant increase in TUG performance timing in the dual-task condition, the increased time of 0.35s is substantially lower than the minimal detectable change that is on the order of seconds not sub-seconds (e.g., 1s in individuals with knee osteoarthritis (39) and 3 second in stroke survivors (40)). The lack of meaningful change in TUG time or counting performance suggests that our young adult population was able to successfully focus on the counting tasks enough to keep their performance consistent without compromising TUG performance.

Although motor module number and composition did not change in the presence of a cognitive distraction, motor module activation became more consistent. This result is in contrast with our hypothesis that activation would become more variable when cognitively distracted. Our finding that motor module activations became more consistent when performing the TUG test with a cognitive distraction could mean that subjects allowed their movements to become more automatic while they focused on the counting task, despite instructions to pay equal attention to both counting and TUG performance. Movements like walking require both automatic and executive control, but healthy adults rely on more automatic control than other populations. In populations that use less automatic control for walking, such as older adults, walking and cognitive tasks compete for executive control resources, impeding performance in both tasks (41). However, healthy young adults likely have enough automaticity and processing capacity to devote attention to the cognitive task while relying on automatic control to perform the TUG test. Our results of increased recruitment consistency are also in agreement with recent work demonstrating increased dynamic stability of motor modules under dual task conditions without corresponding effects on center of mass stability (in anterior/posterior or mediolateral directions (42)), suggesting an adjustment by the nervous system to prioritize stability during cognitive distractions.

Alternatively, the increased activation consistency could be related to the instructions, order of tasks, and/or difficulty of the cognitive task. TUGC trials were always performed second, and subjects may have been more confident paying less attention to their movements than if TUGC had occurred first. Additionally, subjects may not pay much attention to their initial TUG performance but become more focused during TUGC because of the instructions given. For the normal TUG test, subjects are given no instructions about their focus, and may allow their minds to wander during this repetitive and unchallenging task. During TUGC, they are told to pay equal attention to both the counting and TUG and may therefore give the TUG performance more attention than they had previously, leading to more consistent motor module activations. Finally, it is also possible that our findings are influenced by the difficulty of the cognitive task. In particular, the serial subtraction by threes may have been too easy for our young adult population. Decker and colleagues demonstrated a U-shaped relationship between cognitive demand and gait control (measured through step length and width variabilities (43)); more changes in motor module activations could emerge with more difficult dual task conditions.

Though the underlying reasons for the change in motor module activations in the presence of cognitive distraction remain unclear, our results do suggest that cognitive distraction can impact motor module recruitment. Careful follow up studies could clarify the responses by incorporating a variety of cognitive distractions and controlling for practice effects. Understanding how cognitive distractions impact motor module recruitment and activation would provide further insight into the underlying neuromuscular control mechanisms in both healthy and balance impaired populations who may be more affected by cognitive dual tasking.

## 5. Conclusions

Our results support the hypothesis that healthy young adults recruit from a “library” of motor modules to meet the multi-tasks demands of daily life. Specifically, we found that a small number of common motor modules were recruited during walking, turning, and chair transfers and that their structure was robust to cognitive distraction. Achieving different mechanical and cognitive demands were accomplished through changes in motor module activation. This work is the first step towards a full characterization of motor module recruitment patterns in healthy adults across a wide range of daily life tasks. Our results provide a basis for interpreting the effects of motor module changes on mobility and fall risk during daily life that occur in populations with neural or musculoskeletal injuries.

## List of Abbreviations

ADD: adductor magnus
BFLH: biceps femoris long head
C: a # modules x time matrix of motor module activation coefficients, from EMG=W x C + ε
EMG: electromyography
ε: EMG reconstruction error
GMAX: gluteus maximus
GMED: gluteus medius
LGAS: lateral gastrocnemius
MGAS: medial gastrocnemius
PERO: peroneus
RFEM: rectus femoris
SOL: soleus
TA: tibialis anterior
TFL: tensor fasciae latae
TUG: Timed-Up-and-Go Test
TUGC: Timed-Up-and-Go Test with cognitive dual-task
VLAT: vastus lateralis
W: a # muscles x # motor modules matrix of motor modules, from EMG=W x C + ε

