## Supplementary Material for "Young Adults Recruit Similar Motor Modules Across Walking, Turning, and Chair Transfers"

### 1. TUG Subtask Segmentation Example

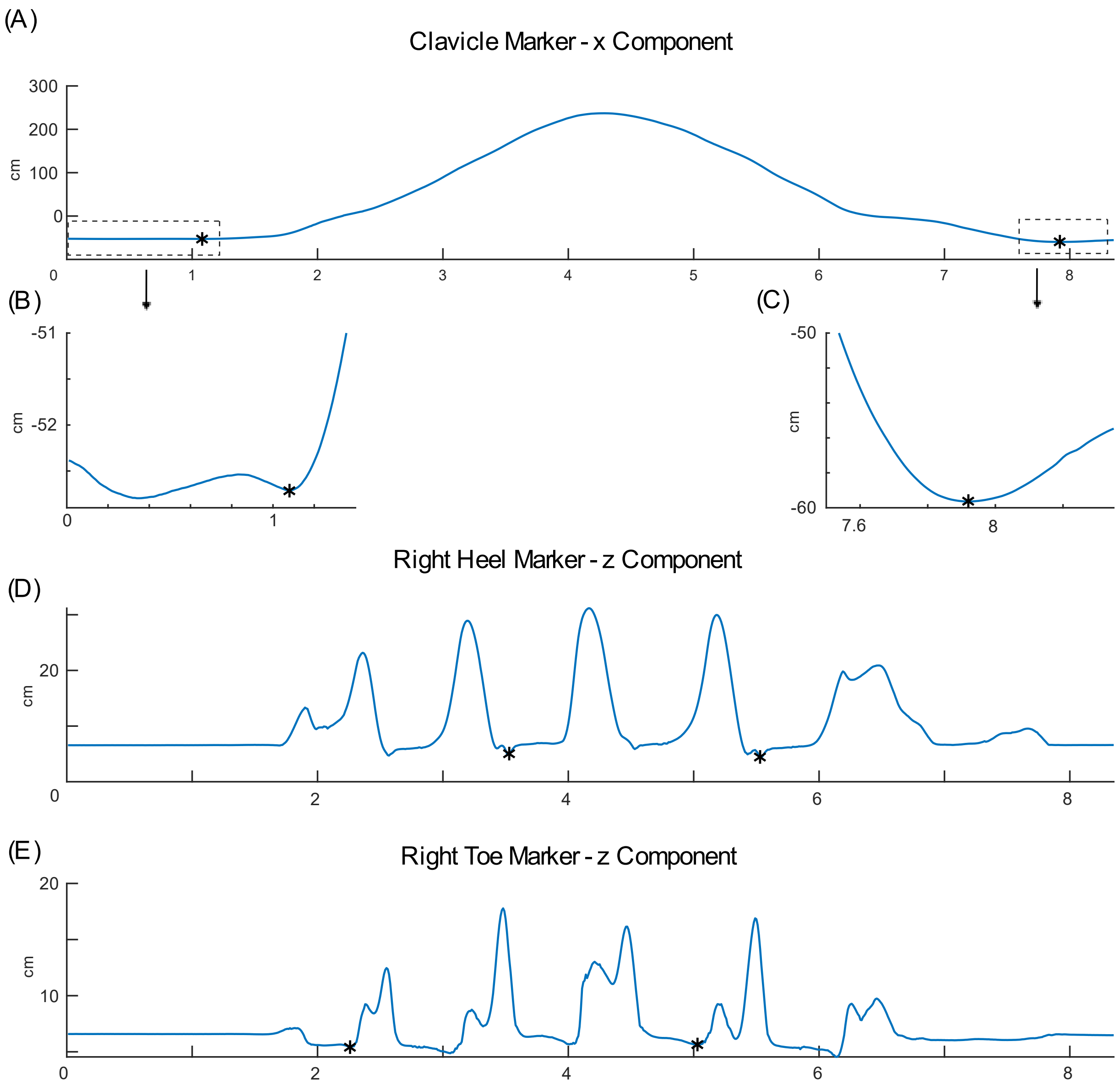


**Figure 1: Example of TUG subtask segmentation in a representative subject.** (A) Clavicle marker in the x-direction, used to identify TUG start and stop. (B) Toe markers in the z-direction, used to identify Walk 1 start and Turn start. (C) Heel markers in the z-direction, used to identify Turn stop and Walk 2 stop. Also see Table 1 in the manuscript.

### 2. Normalization of Motor Module Activations

Each trial was normalized to be the same number of points (1024) and such that the number of data points in Sit-to-Stand, Walk-Turn-Walk, and Stand-to-Sit was consistent. These values were determined by pooling the subtask proportions (subtask time / full TUG time) across all subjects and trials in the single task TUG test, averaging them for each subtask, and rounding to the nearest integer. This yielded

- Sit-to-Stand: 15%
- Walk 1: 17%
- Turn: 25%
- Walk 2: 10%
- Stand-to-Sit: 33%

The Turn and the two Walk subtasks were combined to avoid introducing any experimenter bias from which steps were selected as the beginning and end of the turn. As described above, TUG subtasks were manually identified, so even though subjects were generally consistent with the number of steps, a particular step could be classified as turning or walking in different trials depending on the subject's orientation. Motor modules activations were then normalized as follows:

- Sit to stand - 15% - 154 points
- Walk-Turn-Walk - 52% - 532 points
- Stand-to-Sit - 33% - 338 points

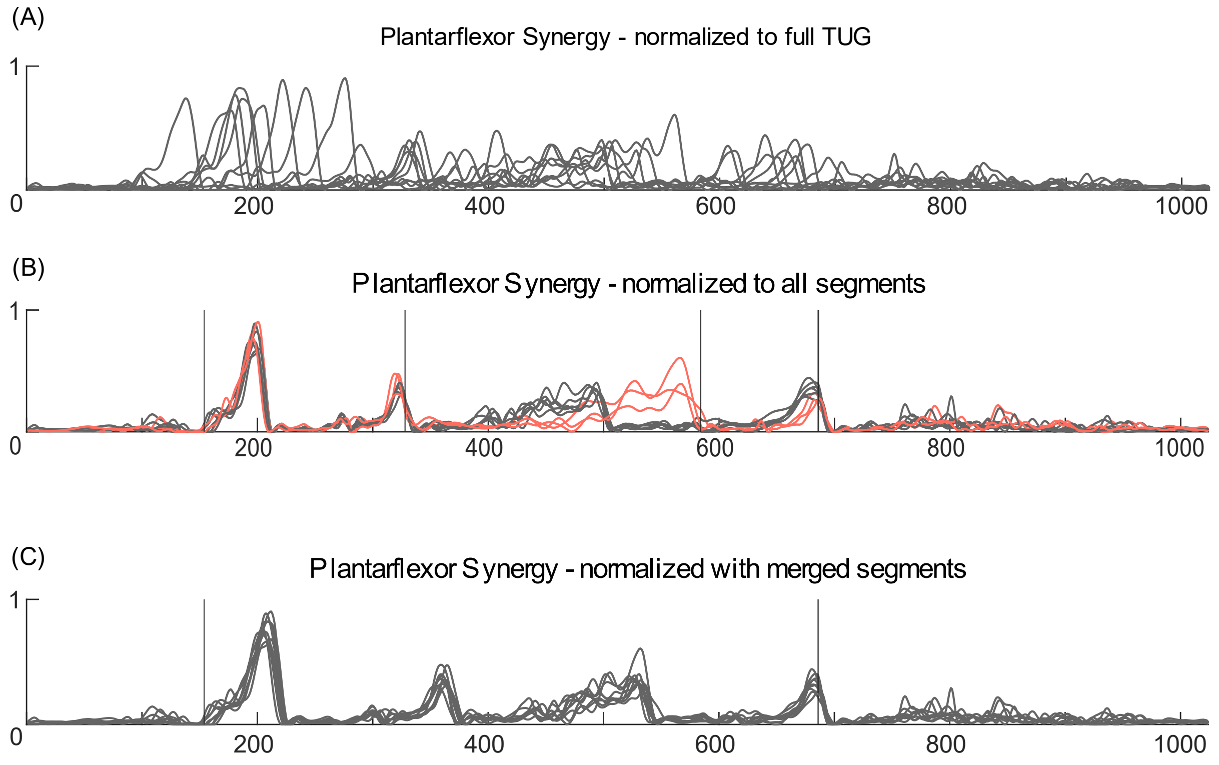


**Figure 2: Example of normalization in a single motor module from a representative subject in the cognitive dual-task TUG test.** (A) Motor module activation profiles normalized to the

full TUG test only. Here the only defined points are the beginning and end of the TUG test. (B) Activation profiles with every subtask normalized. A couple different curve profiles can be seen that could be due to the turn identification, colored in orange. Vertical lines indicate the divisions between TUG subtasks. (C) Activation profiles normalized to three sections (Sit-to-Stand, Walk-Turn-Walk, and Stand-to-Sit), as used in the manuscript. Here, the peaks for each step are more aligned, allowing for better RMSE analysis.

3. Kinematic Strategy Separation

Trials were classified based on "kinematic strategy", defined as the sequence of the first step leg, turn direction, and stand-to-sit turn direction (e.g., RRR for right first step, right turn, stand to sit right turn). The shapes of the activation curves depended on kinematic strategy; for example, the first peak in the plantarflexor synergy would depend on which foot was used to step off. We separated kinematic strategies in this way to avoid artificially analyzing variation in motor module recruitment that was purely due to such kinematic strategy differences.

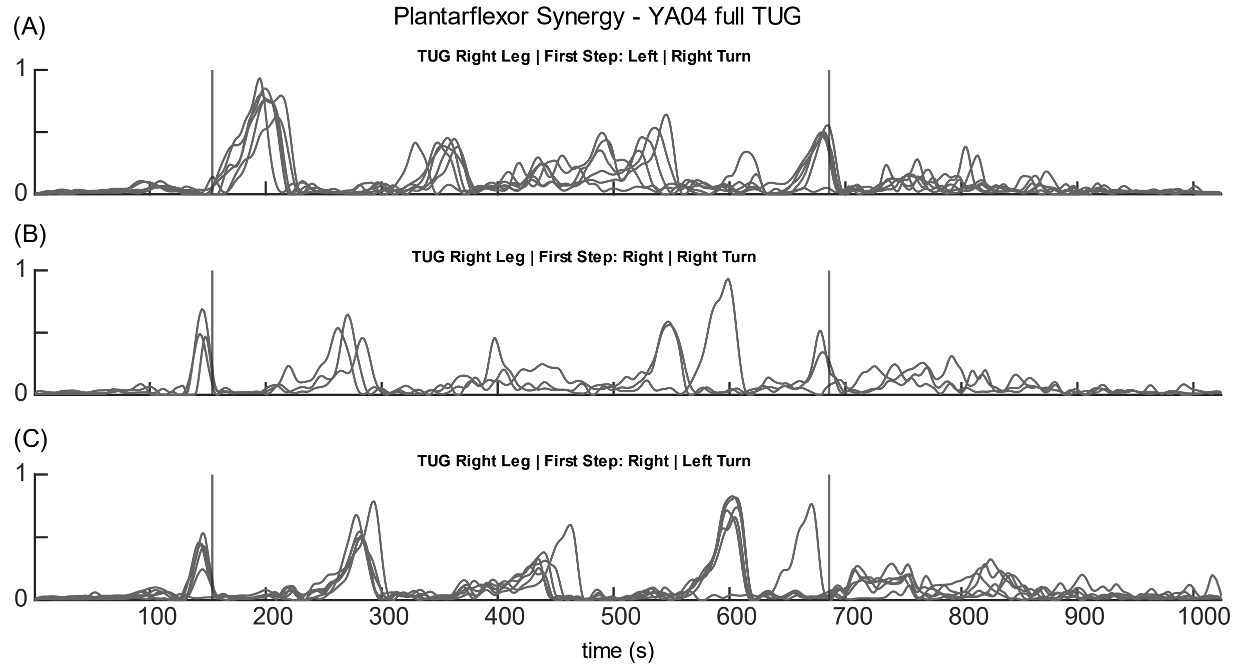


**Figure 3: Example of activation profiles from two motor modules in three kinematic strategies.** Activations from the kinematic sequences LRR (A), RRR (B), and RLR (C).

Because the first leg step-off and stand-to-sit turn direction were not enforced, some subjects varied their kinematic strategies, and did not always use the same ones in TUG and TUGC. Table 2 contains the number of trials in which the subjects used each kinematic strategy. For example, YA23 used 3 kinematic strategies in both TUG and TUGC, but only two (RRR and RLL) were used in both conditions. So that only like performances from TUG and TUGC were compared, we only included trials that had the same kinematic sequences in both TUG and TUGC in the RMSE analysis.

**Table 1: Kinematic Strategies and Number of Trials.** This table contains the number of trials per kinematic sequence for each subject, in the single-task TUG test (TUG) and the dual-task TUG test (TUGC). Each of the 8 kinematic sequences consists of a first step leg, turn direction, and stand-to-sit turn direction. For example, the sequence 'RRR' indicates the subject stepped off with their right foot and turned to the right for both turns; the sequence LRL indicates step off with the left foot, a right turn around the cone, and a left turn before sitting down. Table cells contain either the number of trials that the subject used that sequence, or a dash for sequences that were unused. Colored cells indicate sequences that were used in both TUG and TUGC and were therefore included in the RMSE analysis.

| | TUG | | | | | | | | TUGC | | | | | | | |
|---|---|---|---|---|---|---|---|---|---|---|---|---|---|---|---|---|
| | RRR | LRR | RLR | LLR | RRL | LRL | RLL | LLL | RRR | LRR | RLR | LLR | RRL | LRL | RLL | LLL |
| YA04 | 3 | 6 | 6 | - | - | - | - | - | - | 9 | - | 10 | - | - | - | - |
| YA08 | - | - | - | - | 8 | - | - | 8 | 4 | - | - | 4 | 5 | - | - | 6 |
| YA10 | 8 | - | - | 11 | - | - | - | - | 5 | 1 | - | 11 | - | - | - | - |
| YA11 | 5 | - | 10 | - | - | - | - | - | 10 | - | - | - | - | - | 10 | - |
| YA12 | - | 10 | - | 10 | - | - | - | - | - | 10 | - | 10 | - | - | - | - |
| YA14 | 9 | - | - | - | - | - | 7 | - | - | - | - | - | - | - | 8 | - |
| YA15 | 10 | - | 10 | - | - | - | - | - | 11 | - | 4 | - | - | - | 7 | - |
| YA16 | 10 | - | 7 | 3 | - | - | - | - | 10 | - | 9 | - | - | - | - | - |
| YA18 | 11 | - | - | 1 | - | - | - | 9 | 13 | - | - | - | - | - | 1 | 10 |
| YA19 | 10 | - | 10 | - | - | - | - | - | 10 | - | 10 | - | - | - | - | - |
| YA21 | 10 | 1 | 4 | 5 | - | - | - | - | - | - | 10 | - | 10 | - | - | - |
| YA22 | - | - | - | - | 10 | - | 9 | - | - | - | - | - | 10 | 1 | 10 | - |
| YA23 | 9 | - | - | - | 1 | - | 9 | - | 11 | - | 5 | - | - | - | 7 | - |

4. Detailed Results

**Table 2: Number of motor modules recruited across all subjects and subtasks.**

| | | TUG | | TUGC | | # shared | | % shared | |
|---|---|---|---|---|---|---|---|---|---|
| subject | subtask | L | R | L | R | L | R | L | R |
| YA04 | full | 4 | 4 | 4 | 4 | 4 | 4 | 1 | 1 |
| | Sit-to-Stand | 3 | 3 | 3 | 3 | 3 | 2 | 1 | 0.5 |

| | | | | | | | | | |
|---|---|---|---|---|---|---|---|---|---|
| | Walk | 4 | 3 | 4 | 3 | 4 | 1 | 1 | 0.2 |
| | Turn Left | 4 | 3 | 4 | 4 | 4 | 3 | 1 | 0.75 |
| | Turn Right | 4 | 5 | 3 | 4 | 3 | 3 | 0.75 | 0.5 |
| | Stand-to-Sit Right | 5 | 4 | 4 | 5 | 4 | 4 | 0.8 | 0.8 |
| YA08 | full | 5 | 5 | 5 | 5 | 5 | 5 | 1 | 1 |
| | Sit-to-Stand | 5 | 4 | 4 | 4 | 3 | 3 | 0.5 | 0.6 |
| | Walk | 5 | 5 | 5 | 5 | 5 | 5 | 1 | 1 |
| | Turn Left | 5 | 4 | 4 | 4 | 4 | 4 | 0.8 | 1 |
| | Turn Right | 5 | 5 | 4 | 5 | 4 | 5 | 0.8 | 1 |
| | Stand-to-Sit Left | 5 | 5 | 6 | 5 | 5 | 5 | 0.83 | 1 |
| | Stand-to-Sit Right | n/a | n/a | 6 | 6 | n/a | n/a | n/a | n/a |
| YA10 | full | 5 | 5 | 4 | 4 | 4 | 4 | 0.8 | 0.8 |
| | Sit-to-Stand | 4 | 3 | 4 | 3 | 4 | 2 | 1 | 0.5 |
| | Walk | 4 | 4 | 4 | 4 | 4 | 4 | 1 | 1 |
| | Turn Left | 4 | 4 | 4 | 4 | 2 | 4 | 0.33 | 1 |
| | Turn Right | 3 | 5 | 3 | 5 | 3 | 4 | 1 | 0.67 |
| | Stand-to-Sit Right | 5 | 6 | 4 | 5 | 3 | 5 | 0.5 | 0.83 |
| YA11 | full | 4 | 4 | 5 | 4 | 4 | 4 | 0.8 | 1 |
| | Sit-to-Stand | 3 | 3 | 3 | 3 | 3 | 3 | 1 | 1 |
| | Walk | 3 | 4 | 4 | 4 | 2 | 4 | 0.4 | 1 |
| | Turn Left | 5 | 4 | 5 | 4 | 5 | 4 | 1 | 1 |
| | Turn Right | 5 | 5 | 5 | 5 | 5 | 3 | 1 | 0.43 |
| | Stand-to-Sit Left | n/a | n/a | 5 | 3 | n/a | n/a | n/a | n/a |
| | Stand-to-Sit Right | 4 | 5 | 4 | 5 | 4 | 5 | 1 | 1 |
| YA12 | full | 5 | 5 | 5 | 5 | 5 | 5 | 1 | 1 |
| | Sit-to-Stand | 4 | 3 | 4 | 3 | 3 | 3 | 0.6 | 1 |
| | Walk | 4 | 3 | 4 | 4 | 4 | 2 | 1 | 0.4 |
| | Turn Left | 5 | 4 | 5 | 5 | 3 | 3 | 0.43 | 0.5 |
| | Turn Right | 5 | 5 | 4 | 5 | 4 | 3 | 0.8 | 0.43 |
| | Stand-to-Sit Right | 5 | 5 | 5 | 5 | 5 | 4 | 1 | 0.67 |
| YA14 | full | 5 | 4 | 5 | 3 | 5 | 3 | 1 | 0.75 |
| | Sit-to-Stand | 3 | 3 | 3 | 3 | 2 | 3 | 0.5 | 1 |
| | Walk | 4 | 4 | 4 | 3 | 4 | 3 | 1 | 0.75 |
| | Turn Left | 5 | 3 | 5 | 3 | 5 | 3 | 1 | 1 |
| | Turn Right | 4 | 4 | 4 | 4 | 4 | 4 | 1 | 1 |
| | Stand-to-Sit Left | 5 | 3 | 4 | 3 | 4 | 3 | 0.8 | 1 |
| | Stand-to-Sit Right | 4 | 3 | 4 | 3 | 4 | 3 | 1 | 1 |

| | | | | | | | | | |
|---|---|---|---|---|---|---|---|---|---|
| YA15 | full | 4 | 5 | 5 | 5 | 4 | 5 | 0.8 | 1 |
| | Sit-to-Stand | 3 | 4 | 3 | 4 | 2 | 4 | 0.5 | 1 |
| | Walk | 3 | 4 | 4 | 4 | 2 | 4 | 0.4 | 1 |
| | Turn Left | 4 | 4 | 5 | 4 | 4 | 4 | 0.8 | 1 |
| | Turn Right | 3 | 5 | 4 | 5 | 2 | 5 | 0.4 | 1 |
| | Stand-to-Sit Left | n/a | n/a | 4 | 5 | n/a | n/a | n/a | n/a |
| | Stand-to-Sit Right | 4 | 5 | 5 | 5 | 4 | 5 | 0.8 | 1 |
| YA16 | full | 4 | 5 | 4 | 5 | 4 | 5 | 1 | 1 |
| | Sit-to-Stand | 3 | 3 | 3 | 3 | 2 | 3 | 0.5 | 1 |
| | Walk | 3 | 5 | 3 | 5 | 3 | 5 | 1 | 1 |
| | Turn Left | 5 | 4 | 4 | 4 | 4 | 4 | 0.8 | 1 |
| | Turn Right | 4 | 4 | 4 | 4 | 3 | 3 | 0.6 | 0.6 |
| | Stand-to-Sit Right | 4 | 5 | 4 | 3 | 4 | 3 | 1 | 0.6 |
| YA18 | full | 5 | 5 | 5 | 5 | 4 | 5 | 0.67 | 1 |
| | Sit-to-Stand | 4 | 4 | 4 | 5 | 4 | 4 | 1 | 0.8 |
| | Walk | 4 | 5 | 4 | 4 | 4 | 4 | 1 | 0.8 |
| | Turn Left | 4 | 4 | 4 | 4 | 4 | 3 | 1 | 0.6 |
| | Turn Right | 3 | 5 | 3 | 2 | 2 | 1 | 0.5 | 0.17 |
| | Stand-to-Sit Left | 4 | 5 | 5 | 5 | 3 | 5 | 0.5 | 1 |
| | Stand-to-Sit Right | 5 | 5 | 5 | 6 | 2 | 4 | 0.25 | 0.57 |
| YA19 | full | 4 | 5 | 4 | 5 | 4 | 5 | 1 | 1 |
| | Sit-to-Stand | 2 | 4 | 2 | 4 | 2 | 4 | 1 | 1 |
| | Walk | 3 | 4 | 3 | 5 | 3 | 4 | 1 | 0.8 |
| | Turn Left | 4 | 4 | 4 | 4 | 4 | 4 | 1 | 1 |
| | Turn Right | 4 | 4 | 3 | 5 | 3 | 3 | 0.75 | 0.5 |
| | Stand-to-Sit Right | 5 | 5 | 5 | 5 | 3 | 5 | 0.43 | 1 |
| YA21 | full | 4 | 5 | 5 | 5 | 4 | 5 | 0.8 | 1 |
| | Sit-to-Stand | 3 | 3 | 2 | 3 | 1 | 3 | 0.25 | 1 |
| | Walk | 4 | 5 | 3 | 5 | 2 | 3 | 0.4 | 0.43 |
| | Turn Left | 4 | 5 | 5 | 5 | 3 | 5 | 0.5 | 1 |
| | Turn Right | 4 | 4 | 5 | 4 | 4 | 4 | 0.8 | 1 |
| | Stand-to-Sit Left | n/a | n/a | 6 | 5 | n/a | n/a | n/a | n/a |
| | Stand-to-Sit Right | 5 | 6 | 4 | 6 | 4 | 6 | 0.8 | 1 |
| YA22 | full | 4 | 4 | 4 | 4 | 4 | 4 | 1 | 1 |
| | Sit-to-Stand | 3 | 3 | 3 | 3 | 2 | 3 | 0.5 | 1 |
| | Walk | 4 | 4 | 3 | 4 | 3 | 4 | 0.75 | 1 |
| | Turn Left | 4 | 3 | 5 | 3 | 4 | 3 | 0.8 | 1 |
| | Turn Right | 4 | 4 | 3 | 4 | 3 | 4 | 0.75 | 1 |

| | | | | | | | | | |
|---|---|---|---|---|---|---|---|---|---|
| | Stand-to-Sit Left | 5 | 5 | 4 | 5 | 4 | 5 | 0.8 | 1 |
| YA23 | full | 4 | 4 | 4 | 5 | 4 | 4 | 1 | 0.8 |
| | Sit-to-Stand | 3 | 4 | 2 | 3 | 2 | 3 | 0.67 | 0.75 |
| | Walk | 4 | 5 | 4 | 5 | 3 | 3 | 0.6 | 0.43 |
| | Turn Left | 5 | 4 | 5 | 4 | 5 | 2 | 1 | 0.33 |
| | Turn Right | 4 | 4 | 4 | 4 | 4 | 4 | 1 | 1 |
| | Stand-to-Sit Left | 4 | 4 | 4 | 5 | 3 | 4 | 0.6 | 0.8 |
| | Stand-to-Sit Right | 4 | 4 | 5 | 4 | 3 | 4 | 0.5 | 1 |

**Table 3: Results of t-tests comparing motor module composition in TUG vs TUGC for all TUG subtasks**

| | p-value |
|---|---|
| full | 0.713 |
| Sit-to-Stand | 0.185 |
| Walk | 1 |
| Turn Left | 0.265 |
| Turn Right | 0.161 |
| Stand-to-Sit Left Turn | 0.678 |
| Stand-to-Sit Right Turn | 0.576 |

**Table 4: Clustering Results.**

| | number of clusters | | % generalization | | Avg Cluster Consistency | | % shared with Full TUG Test | |
|---|---|---|---|---|---|---|---|---|
| subject | Left Leg | Right Leg | Left Leg | Right Leg | Left Leg | Right Leg | Left Leg | Right Leg |
| YA04 | 5 | 5 | 90 | 89 | 0.85 | 0.86 | 80 | 80 |
| YA08 | 8 | 5 | 88 | 96 | 0.81 | 0.82 | 63 | 100 |
| YA10 | 5 | 6 | 90 | 86 | 0.77 | 0.83 | 100 | 83 |
| YA11 | 5 | 5 | 90 | 90 | 0.83 | 0.80 | 80 | 80 |
| YA12 | 6 | 5 | 91 | 90 | 0.82 | 0.60 | 83 | 67 |
| YA14 | 5 | 4 | 92 | 95 | 0.82 | 0.84 | 67 | 100 |
| YA15 | 5 | 5 | 88 | 95 | 0.72 | 0.78 | 80 | 100 |
| YA16 | 5 | 7 | 89 | 81 | 0.81 | 0.85 | 80 | 71 |
| YA18 | 6 | 7 | 88 | 89 | 0.82 | 0.77 | 38 | 71 |
| YA19 | 5 | 6 | 83 | 90 | 0.78 | 0.70 | 80 | 83 |
| YA21 | 6 | 6 | 85 | 87 | 0.71 | 0.74 | 67 | 83 |
| YA22 | 5 | 5 | 90 | 89 | 0.83 | 0.79 | 80 | 80 |
| YA23 | 5 | 7 | 92 | 88 | 0.78 | 0.89 | 80 | 57 |
